# A minimal yet flexible likelihood framework to assess correlated evolution

**DOI:** 10.1101/2020.09.04.282954

**Authors:** Abdelkader Behdenna, Maxime Godfroid, Patrice Petot, Joël Pothier, Camille Nous, Amaury Lambert, Guillaume Achaz

## Abstract

An evolutionary process is reflected in the sequence of changes of any trait (e.g. morphological, molecular) through time. Yet, a better understanding of evolution would be procured by characterizing correlated evolution, or when two or more evolutionary processes interact. Many previously developed parametric methods often require significant computing time as they rely on the estimation of many parameters. Here we propose a minimal likelihood framework modelling the joint evolution of two traits on a known phylogenetic tree. The type and strength of correlated evolution is characterized by few parameters tuning mutation rates of each trait and interdependencies between these rates. The framework can be applied to study any discrete trait or character ranging from nucleotide substitution to gain or loss of a biological function. More specifically, it can be used to 1) test for independence between two evolutionary processes, 2) identify the type of interaction between them and 3) estimate parameter values of the most likely model of interaction. In its current implementation, the method takes as input a phylogenetic tree together with mapped discrete evolutionary events on it and then maximizes the likelihood for one or several chosen scenarios. The strengths and limits of the method, as well as its relative power when compared to a few other methods, are assessed using both simulations and data from 16S rRNA sequences in a sample of 54 γ-enterobacteria. We show that even with datasets of fewer than 100 species, the method performs well in parameter estimation and in the selection of evolutionary scenario.

## Introduction

Evolutionary processes are often interdependent at all levels of organization, from molecules (Shindyalov et al., 1994) to ecosystems (Van Valen, 1973). *Correlated evolution* is the lack of independence in the evolution of multiple traits of the same living entity such as an individual, a sequence or a species (Achaz and Dutheil, 2021). Here, we define *trait* as any morphological or life history trait, or any molecular character. In many cases, one mutation in one trait impacts the evolution of other traits. Patterns of correlated evolution naturally emerges when comparing morphological traits (e.g., size and body mass) or when comparing molecular traits (e.g., residues of the same protein). Nevertheless, correlated evolution might also be observed through functional constraints in metabolic or regulation networks, without any physical interactions between the partners (Fraser et al., 2004).

The nature of biological interactions explains the different patterns of correlated evolution. For example, synergistic effects result in positive correlation whereas antagonistic ones cause negative correlations. At the molecular level, genetic interactions are named epistasis (Phillips, 2008), which refers to the influence of a genetic background on the effect of mutations. Multiple mutations may have additive or diminishing fitness effects, and all combinations are summarized in a fitness landscape. A fitness landscape is then the catalog of fitness values of all possible combinations of mutations (Wright, 1932), and its structure is highly predictive of the expected pattern of correlated evolution (Achaz and Dutheil, 2021). Fitness landscapes are an object of still important work nowadays (Achaz et al., 2014; Visser and Krug, 2014; Yi and Dean, 2019).

In this study, we restrict ourselves to the simple case of only two evolutionary processes. Compensated Pathogenic Deviations (CPDs) constitute an illuminating example of correlated evolution. CPDs are instances of the more general Bateson-Dobzhansky-Muller incompatibilities (Bateson, 1909; Dobzhansky, 1934; Muller, 1942; Orr, 1996; Welch, 2004). CPDs are highly deleterious alleles in a focal species but wild type in one or several other species. The deleterious effect of the mutated allele must then be balanced by one or more *compensatory* mutations, assuming that the effect is not simply due to environmental effects. CPDs have been reported for humans (Kondrashov et al., 2002) and insects (Kulathinal et al., 2004), and they can be as common as 10% of the deleterious amino acid substitutions. Depending on the exact nature of the epistasis between the two mutations, different mutational histories will have different likelihoods. If the first mutation to occur is the deleterious one, then the occurrence of the compensatory mutation must follow very quickly or even co-occur with the first one. When the compensatory mutation occurs first, the second one, that is no longer deleterious, can occur with some time lag. Both possible scenarios can be re-expressed in terms of induction of one event onto the other; the induction represents the intensity in which the triggering event favors the other. The inferred strength of the induction will be however different for both scenarios: for example, a strong induction in the first case and only a mild induction in the second one.

Although direct interactions cause patterns of correlated evolution, the reciprocal is not true, for at least two reasons.

First, an observed pattern of correlation can result from indirect interactions between the two focal traits. As always, correlation does not imply causation. For example, two morphological traits can jointly respond to changes of a hidden environmental variable without having direct interaction. Similarly, two residues in proteins can show patterns of correlated evolution because they both interact with a third residue. To overcome this issue, the popular Direct Coupling Analysis (DCA) has been developed, which prevents indirect coupling and allows only pairwise interactions (Weigt et al., 2009; Morcos et al., 2011). In particular, DCA models the abundance of sequences in nature by probabilities of presence in a Potts model. In such a model, the distribution of sequence abundance depends not only on marginal frequencies at each site (local abundance) but also on joint frequencies for all pairs of sites (pairwise epistasis). The underlying assumption is that deleterious combinations of residues should be rare enough such that we do not see them in the wild. The main challenge of DCA is to infer the parameters of a highly parameterized model. Several methods have been developed to infer correctly and efficiently the model parameters but they are computationally demanding (Weigt et al., 2009; Morcos et al., 2011; Baldassi et al., 2014; Ekeberg et al., 2013). The advantage of DCA is the excellent predictive power of amino acid contacts in 3D structures (Marks et al., 2011) in comparison to simple correlation metrics such as Mutual Information (MI) (Chiu and Kolodziejczak, 1991; Martin et al., 2005). Nonetheless, MI methods are decent indicators of interactions and are so fast to compute that they can be measured for all pairs of sites on alignments of complete bacterial genomes (Bitbol, 2018; Pensar et al., 2019).

Second, patterns of apparent correlated evolution can be due to phylogenetic inertia (Harvey and Pagel, 1991). More specifically, species, sequences or individuals cannot be regarded as statistically independent samples because they partially share a history. This shared history is often represented by a phylogenetic tree. Species closer in the tree have traits that are closer in value and this extends to pairs of traits. Hence, the variable phylogenetic proximity between species create patterns of correlation for traits that are not biologically interacting. In this regard, phylogeny can be seen as a hidden variable. For continuous traits, the development of the statistically independent contrasts led to take into account phylogenetic inertia (Felsenstein, 1985). For discrete traits, an extension of the phylogenetic logistic regression for binary dependent variables was recently developed (Ives and Garland, 2010).

In this article, we are interested in quantifying correlated evolution between discrete traits. We will restrict the term *evolutionary process* to describe any process that results in *discrete evolutionary events* on a phylogenetic tree. Discrete events abstract mutations at the molecular level, but more generally can be applied to any change, gain, or loss of any biological function, of any morphological trait or of any life history trait. These events can be mapped on the phylogenetic tree using ancestral character reconstruction (Shindyalov et al., 1994). Once the events are mapped, one can then test for the independence between the evolutionary paths of the two traits and whether both traits tend to show co-variation in the same branches of the tree (Tufféry and Darlu, 2000; Dutheil et al., 2005; Dutheil and Galtier, 2007) or whether one type of event tends to precede the other (Kryazhimskiy et al., 2011; Behdenna et al., 2016). For cases where events are placed (ignoring reconstruction uncertainties), p-values can be computed analytically for any type of correlated evolution (one event precedes the other, both events co-occur, one event prohibits the other, etc.) using matrix formalism (Behdenna et al., 2016).

The null model of independence assumes that evolutionary events follow a Poisson process, such that they occur uniformly on the tree. More generally, evolutionary processes can be modeled by time-continuous Markov processes (Felsenstein, 1981). It is then possible to generalize the model for two evolutionary processes that explicitly depend on one another (Pagel, 1994; Milligan, 1994). The model of independence corresponding to a sub-space of parameter values of the general model makes it possible to test whether the two evolutionary processes are independent (or not) using likelihood-ratio tests (LRT). Based on this idea, multiple likelihood-based approaches were proposed, although they are generally computationally demanding (Pagel, 1994; Milligan, 1994; Schoniger and von Haeseler, 1994; Tillier and Collins, 1995; Pollock et al., 1999; Baum and Donoghue, 2001; Pagel and Meade, 2006; Yeang et al., 2007; Dib et al., 2014). The methods involve exploring or maximizing over a likelihood surface that has as many dimensions as the number of parameters. For two binary traits, there are 4 possible states for the two traits (00, 01, 10, 11). When the two processes are independent, there are four transition rates (two for each trait in both directions) but there is a 4-by-4 rate matrix describing the complete process. The rate matrix is defined by 12 parameters, or 8 when double mutations are forbidden. More complex models, like 2 sites with 20 amino acids each, depend on more parameters (on the order of a few hundred). Consequently, these methods need larger datasets and expensive computation time.

Here, we propose a likelihood framework that is based on a minimal model of correlated evolution. The framework generates a series of nested models with 2 to 8 parameters that correspond to the mutation rates of each process and interactions between them. We show that these parameters are core values that characterize patterns of correlated evolution. The current implementation of the method assumes that the tree is correctly inferred and that the events are correctly placed on the tree. Using maximum likelihood, the method can (i) test for independence between two evolutionary processes, (ii) find the most likely type of correlated evolution between them (obligate or preferential sequential order, reciprocal synergy, incompatibility, etc.) and (iii) estimate the strength of the interaction.

## Materials and Methods

### Model

#### A minimal model of correlated evolution

The following model details the joint distribution of two processes describing sequences of evolutionary events *E_1_* and *E_2_* respectively, on a given tree *T*.

We assume that events *E_i_, i* ∈ {1,2} occur on the branches of tree *T* according to a Poisson process with intensity *m_i_* hereafter called *occurrence rate.* The originality of our model is that the value of *m_i_* can change as a function of the *realization of events E_i_* and *E_j_* on the tree, as we now describe.

For each event *E_i_*, its occurrence rate *m_i_* can take 4 values *μ_i_*, 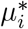, *v_i_* or 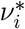. The idea is that the trait modeled by process *i* can take two values, e.g. A or a for process 1 and B or b for process 2. Additionally, traits can be in a basal state or in an excited state, resulting in four states, e.g. for process 1: *A, A*, a, a*,* as depicted in Figure 1a:

- The two basal rates *μ_i_* and *v_i_* are meant to represent natural rates of occurrence of the event *E_i_*, when trait *i* is in its basal state (non-starred), e.g. for process 1: A mutates to *a* at rate *μ*_1_ and *a* mutates to A at rate *v*_1_.
- The starred rates 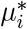 and 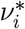 are meant to model excited rates of occurrence of the event *E_i_*, when trait *i* is in its excited state (starred), e.g. for process 1: *A*\* mutates to *a* at rate 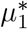 and *a\** mutates to *A* at rate 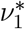. Note that a trait in its excited state returns to a basal state upon mutation.
- Last, a trait *i* can jump from the basal state to the excited state, e.g. for process 1, from *A* to *A*\* or from *a* to *a*\*. These events are induced by occurrences of the other type of event, here *E_j_* with *j* ≠ *i*.

**Figure 1:**
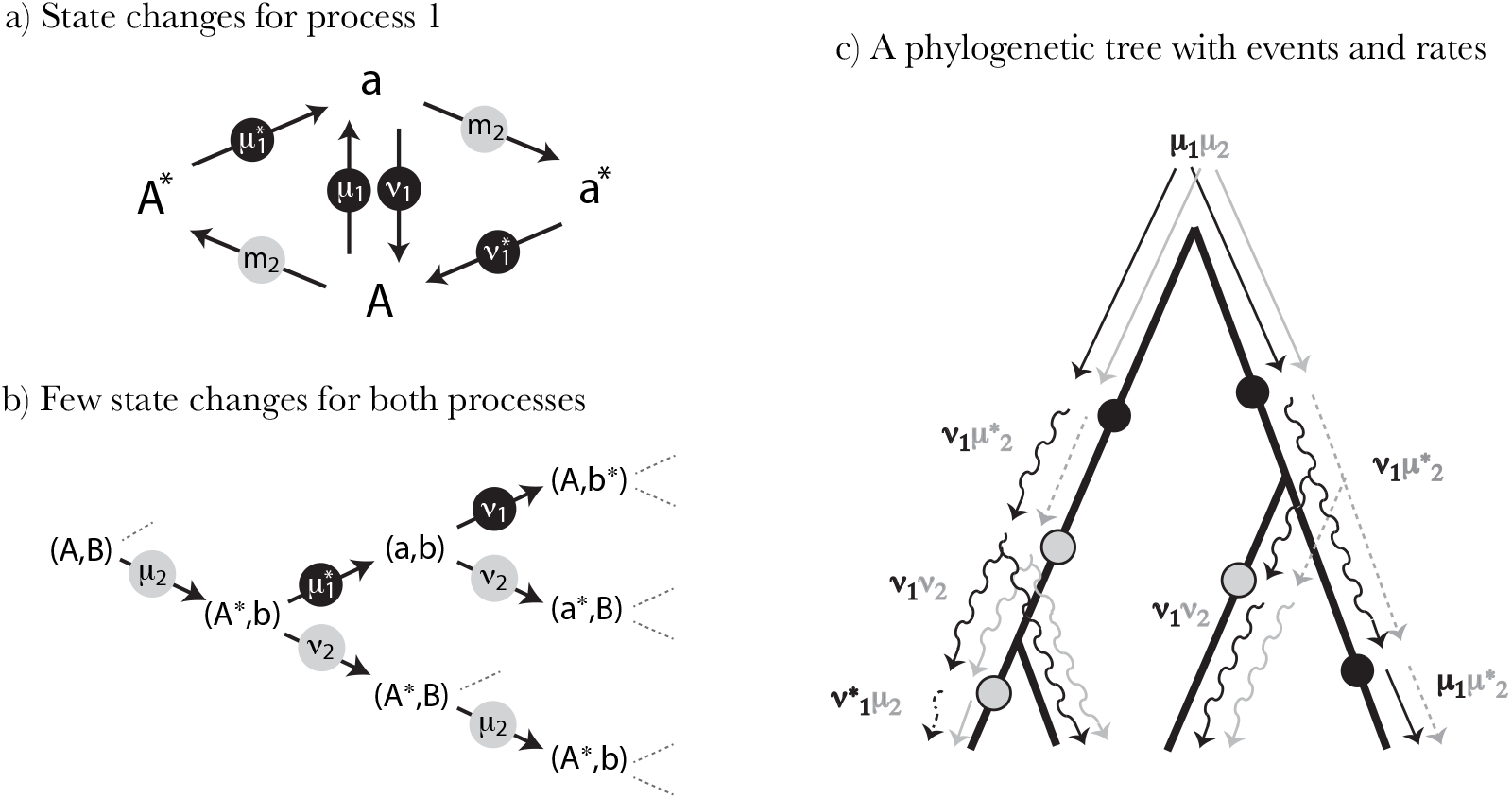
A minimal model of correlated evolution between two processes. Process 1 can be in state *A* or *a* and process 2 can be in state *B* or *b.* Black disks correspond to *E*_1_ events and grey disks correspond to *E*_2_ events. (a) State changes for process 1. Illustration showing the switches between the hidden states of process 1 and the associated occurrence rates indicated within the disks. The occurrence rates *m*_2_ can be either *μ*_2_ or *v*_2_. (b) Subset of hidden states and occurrence rate switches for the two processes. (c) Illustration showing the occurrence rate switches on a mock phylogeny. At the root, the initial rates are μi and *μ*_2_ and the hidden state is *A, B*.

Alternatively, the model can be fully specified by characterizing states with the *rates at which the next event will occur* and thanks to the following two rules (Figure 1):

1. When an *E_j_* event occurs at a non-starred rate (the process *j* was either at *μ_j_* or *v_j_*), the *m_i_* rate of process *i* simultaneously switches to (or remains in) its corresponding starred rate: from *μ_i_* (or 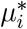) to 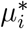, and similarly, from *v_i_* (or 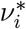) to 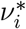.
2. When an *E_i_* event occurs, *m_i_* switches from *μ_i_* (or 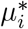) to *v_i_*, and conversely, from *v_i_* (or 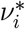) to *μ_i_*, simultaneously losing its star if any.

The ratio 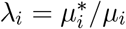 (potentially also 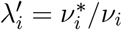) can be thought of as an *induction* factor which measures the influence of an event on another. There is no order requirement between the rates: if *λ_i_* > 1, the induction is positive (i.e. the intensity of the process *i* is increased after an event *E_j_*) whereas it is negative when *λ_i_* < 1. We assume that the interaction only concerns the next induced event, as it models the impact of an *E_j_* event on the rate of occurrence of the next *E_i_* event (and *vice-versa*). Framed in terms of compensatory events, the second event can be seen as compensating for the need created by the first one. In this sense, rule 2 also consumes the induction, as starred rates switch back to non-starred ones. It is noteworthy to mention that, in this model, a second subsequent occurrence of *E_j_* will not set back the rate of *i* to a basal one (the need remains); only an *E_i_* event will do it. Finally, rule 2 also sets the back and forth transitions between the *μ* and *v* rates.

This framework can be used to evaluate many relevant biological scenarios, some of which will be detailed further in the methods.

### The likelihood framework

Let *E*_1_ and *E*_2_ be two types of events whose occurrences are distributed on a tree T, for which the topology and branch lengths are known. We compute the likelihood function as the probability of the positions of the occurrences of both events on T, under the model previously described, conditioned on a set of parameters (*μ_i_*, 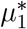,*v_i_*,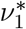, *μ*_2_,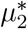, *v*_2_, 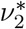). As described below, we can also estimate the rates of the process at the root as well as the order of all events in the tree.

In our current implementation, we assume that no more than one event of each type can occur on a single branch. In the limit of small occurrence rates, this is an excellent approximation. The derivation of the likelihood for more than one event of each type on a branch becomes cumbersome otherwise. As a cautious measure, we calculate the probability that more than one event can occur on the longest branch using the highest basal mutation rate given by the ML estimates. The user is issued a warning whenever the probability is higher than 0.05.

#### The likelihood on a single branch

The likelihood of a branch 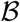 of the tree depends on (i) the branch length, (ii) the occurrence rates at the beginning of the branch and (iii) which event(s) occur on the branch. Considering an initial state of rates (*m*_1_, *m*_2_) and the possible occurrences of *E_1_* and/or *E*_2_ on the branch, we can determine the final state of rates of the current branch, which will be used as the initial rate for the daughter branches, when they exist. All transitions are deterministic and provided in the Appendix. It is noteworthy to mention that, in the model described here, both processes can never be at the same time in a starred rate.

The likelihood of the branch 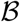 is calculated through the following cases (formula and derivations are detailed in the Appendix): (1) no event occurred on the branch, (2) one event of type *i* occurred on the branch, for *i* ∈ [1, 2] and finally (3) one event of each type (*i* followed by *j* or *j* followed by *i*) occurred on the branch. In the third case, where *E*_1_ and *E*_2_ occurred on 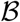, we cannot assume any order between them, as both orders (*E*_1_ → *E*_2_) and (*E*_2_ → *E*_1_) are possible *a priori*; furthermore, the end rate of both processes after the branch depends on this order. We solve this issue by computing the likelihood of the branch for both possible orders (see below).

#### The likelihood on the whole tree

The initial rate state at the root is either (*μ*_1_,*μ*_2_), (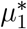,*μ*_2_), or (*μ*_1_,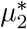). All other pairs are either impossible (e.g. 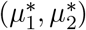) or are identical to a swap between *μ* with *v*. Indeed, *μ* and *v* can be swapped without any loss of generality (e.g. (*μ*_1_,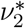) is the same as (*μ*_1_,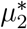) in the model). This remark is useful to limit the computing time, and has no consequence, since the *μ* rates and the *v* rates play a completely symmetric role in our model. We calculate separately the likelihoods considering the 3 possible initial states, using the following algorithm:

- We know the initial rates of both processes on a branch (i.e., inherited from the ancestor branch or given by initialization at the root edge).
- We verify the occurrence of zero, one or both events on the branch concerned and calculate its likelihood and the rates of both processes at the end of the branch. If the branch has both types of events, we calculate separately the likelihood for the two sub-cases and propagate them in parallel with their two respective final rates.
- If the branch is not terminal, we apply this algorithm to its daughter branches, transmitting the final rate. In the case of both events in the branch, we transmit both possible final rates separately to two different paths of the recursion.

The likelihood of the whole tree is the product of the likelihoods of its branches. The number of full-tree likelihoods that must be computed scales with 2^b_2_^, where b_2_ is the number of branches with both types of events. These two-event branches are the main limitation of the method in terms of computation time.

#### Estimating the maximum likelihood

The set of eight parameters (*μ*_1_,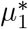, v_1_, 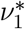,*μ*_2_, 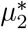, *v*_2_, 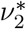) maximizing the likelihood describes the most likely scenario leading to the observed joint distribution of the occurrences of *E*_1_ and *E*_2_ on the tree *T* under our model of correlated evolution. The maximum likelihood (ML) is searched by a dedicated implementation based on a mix of (a) sequential unidimensional gradient ascent (each dimension is optimized successively), (b) multidimensional gradient ascent, both using golden section search and (c) a Newton-Raphson algorithm when possible (when the curvature is negative). The implementation has been optimized for the full model but also for each nested submodels.

Each possible sequence of ordered events from the root to the leaves of the tree is evaluated and the most likely is returned. As an option, we also estimate which one of the three possible initial rates is more likely (maximizing the likelihood for each of them independently); by default we assume that it is (*μ*_1_,*μ*_2_). As a result, the method returns the ordered events, the initial rates (optionally) and the set of parameters maximizing the likelihood.

#### A selection of scenarios

Having two rate categories (*μ/μ*\* and *v/v*\*) for the same process allows us to cover a wider range of scenarios. In this framework, constraining one or more parameters to a fixed value, or to be equal, can define a nested sequence of models, each being a particular case of a more general one (the most general having 8 free parameters). The likelihood can then be maximized for submodels, reducing the parameter space and shortening computation time: for example, setting 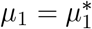 and 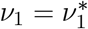 is equivalent to assuming that the occurrence rate of *E*_1_ is not influenced by the occurrences of *E*_2_. Cases of particular interest involve submodels without correlated evolution. There are at least two such models in our framework. The most simplistic one has only two parameters: *μ*_1_ and *μ*_2_ (or even a single parameter *μ* = *μ*_1_ = *μ*_2_). A more flexible version of independence allows for intrinsic rate changes and is modeled by four parameters: μ1, μ2, v1 and v2, as we detail below.

##### Scenario of independence (*H*_0_)

In this model, each event has a single occurrence rate. Indeed, when two processes are independent, their intensities are constant regardless of the position of the occurrences of each type of event on the tree. The model therefore reduces to two parameters *μ*_1_ and *μ*_2_ (assuming that the *v* rates equal the *μ* rates):

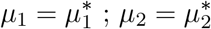

The intensities of both processes are constant in the whole tree. When the two processes are independent but have more than one rate, a more relevant approach is to consider a model with four parameters, where starred rates are equal to non-starred ones but where *μ* rates differ from *v* rates.

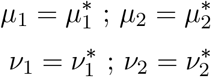

Interestingly, one can also test the relevance of intrinsic rate variations by comparing these two models of independence. Then, one can test for correlated evolution by measuring (and statistically testing) the likelihood improvement with extra parameters, letting the starred rates being different from the basal ones. Some scenarios that our model can calculate are scenarios of prohibition, asymmetric induction and reciprocal induction.

##### Scenario of prohibition

This scenario models the case where *E*_2_ events are not allowed unless they occur after an *E*_1_ event (i.e., *E*_1_ triggers the appearance of E2). This would be the case of a strongly deleterious mutation that cannot occur without being compensated upstream. If we assume a single rate (*μ* = *v*), the model contains 3 parameters

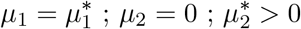

Again, by letting *v* rates differ from *μ* rates, one can also expand the model to include two rates for each process.

##### Scenario of asymmetric induction

*E*_1_ events have a constant rate while the rate of *E*_2_ is increased after an *E*_1_ event has occurred. This scenario models, for example, a compensatory mutation that follows the fixation of a slightly deleterious mutation. More generally, this 3-parameter model describes the process by which direct compensatory events occur.

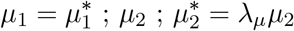

If one wants to include a second rate for both processes, it is possible to add two or three parameters (e.g. 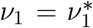; *v*_2_; 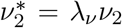), depending on whether one wants to have *λ_μ_* = *λ_ν_* or not. Interestingly, unless one is specifically interested in positive induction, there is no need to restrict *λ*(*_μ,ν_*) > 1, as *λ*(_*μ,ν*_) < 1 would result in a scenario where occurrences of *E*_1_ would slow down the second evolutionary process.

##### Scenario of reciprocal induction

Here, we consider a symmetric interaction between both processes. *E*_1_ events enhance the occurrence rate of *E*_2_ events and, reciprocally, *E*_2_ events enhance the rate of occurrence of *E*_1_ events. This scenario models the reciprocal sign epistasis, where two mutations are beneficial only in the double mutant (Weinreich et al., 2005; Poelwijk et al., 2007).

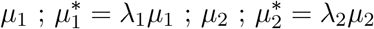

Several variants of this scenario also exist depending on whether *λ*_1_ = *λ*_2_, whether the *λ* are restricted to be larger or smaller than one, and finally whether the *μ* rates differ from the *v* rates.

We can then compare pairs of nested models using LRTs (Neyman and Pearson, 1933). For example, when considering only two nested models, we can use the fact that twice the difference of their maximum log-likelihoods approximately follows a *χ*^2^ distribution with degrees of freedom (df) equal to the difference in numbers of parameters: 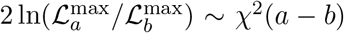. Then by comparing 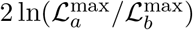 to the quantiles of *χ*^2^(*a* — *b*), we can decide to reject or not the model with more parameters. We have not yet come up with a generalist decision tree using a standard series LRT with a growing number of parameters. We therefore recommend to compute the ML for several scenarios and determine which parameters significantly improve the likelihood (as computation time is short enough). This exploration of the different scenarios that are embedded in the general framework (that has at most eight parameters) will provide an excellent guide to select which scenario is the most likely for the two evolutionary processes under study.

## Simulations

To assess the performance of the method, we simulated different scenarios on a perfectly symmetric phylogenetic tree with six synchronized series of bifurcations and 64 leaves. All branches have the same length.

### Estimating the occurrence rates

We first tested the accuracy of the method to retrieve occurrence rates that have been arbitrarily selected for a series of simulations. We simulated 10, 000 replicates of a scenario where *μ*_1_ = 5, *μ*_2_ = 4, 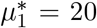 and 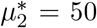 (all *v* were equal to the *μ*). As rates are rescaled by total tree length, *μ*_1_ = 5 implies that there are on average five occurrences of *E*_1_ when the process is at its basal rate (*i.e.*, not counting the ones that are triggered by occurrences of *E*_2_). We applied the ML method to each simulated replicate generating a mapping of events *E*_1_ and *E*_2_ on the tree, in order to estimate thefour rates (the *v* were equal to the *μ* in these simulations). When more than one event of a kind occurred in the branch, only one was kept.

### Power to detect scenarios of induction

To assess the power of LRTs to detect an induction of *E*_1_ on *E*_2_, we simulated two scenarios of induction.

First, we simulated a scenario of asymmetric induction of *E*_1_ on *E*_2_ with three parameters: *μ*_1_ = *μ*_2_ = 5 and the induction rate *λ*_2_ increasing from 1 to 1000. All *v* values are equal to the corresponding *μ*. We then estimate the ML values of three models: an independence model with two parameters (*μ*_1_,*μ*_2_; *H*_0_), the simulated model with three parameters (*H*_1_) and an overparametrized model with inductions in both directions (*μ*_1_, *μ*_2_, 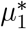, 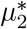; *H*_2_).

The second scenario is a reciprocal induction with three parameters: *μ*_1_ = *μ*_2_ = 5 and *λ*_1_ = *λ*_2_ increasing from 1 to 1000. We estimate the ML values of three models: the same independence model as above (*H*_0_), the simulated model with three parameters (*H*_1_) and an overparametrized model with inductions in both directions (*H*_2_).

In both simulations setup, we compute the LRT of *H*_0_ vs *H*_1_ *(*LRT*_23_,* one degree of freedom) and *H*_2_ vs *H*_1_ (*LRT*_24_, two degrees of freedom). The p-values of significance are computed assuming a *χ*^2^ distribution of twice the logarithm of the ML ratios for their respective degrees of freedom. We compare the results of the LRT to our previous method based on counts of co-occurrences (epics-Id) and on counts of co-occurrences and subsequent occurrences (epicsS+Id) (Behdenna et al., 2016). Additionally, we added a standard analysis for the detection of correlated evolution implemented in BayesTraits Discrete (BTDiscrete) (Pagel, 1994). The last method computes the LRT between a scenario of independence with four parameters and a scenario of dependence with eight parameters. The p-value is thus calculated assuming a *χ*^2^ distribution with four degrees of freedom. We retain all pairs that are found to reject significantly the scenario of independence with a risk of five percent.

### Power to detect induction and/or irreversible loss

We next designed three different scenarios to represent a mixture of intrinsic rate changes and interactions between the events. More specifically, we designed the following scenarios, such that in every case *E*_1_ tends to precede *E*_2_:

1. **Induction**: E_1_ events favors *E*_2_ events (*λ*_2_ > 1), while *E*_2_ events slow down E_1_ events (*λ*_1_ < 1). This is a special case of asymmetric induction of E_1_ on *E*_2_.
2. **Loss**: *E*_1_ events have a higher rate of occurrence (*μ*_1_ > *μ*_2_) and both events can occur only once (*v*_1_ = *v*_2_ = 0). In this scenario, there is no interaction between the two processes. Therefore, the sequential pattern (*E*_1_ precedes *E*_2_) is only due to their respective intrinsic rates. A biological example would be the irreversible loss of two functions, one being lost faster.
3. **Induction+Loss**: this scenario is a combination of irreversible loss (*v*_1_ = *v*_2_ = 0) plus asymmetric induction (*λ*_1_ < 1, *λ*_2_ > 1).

The exact rates of the simulations are indicated in Table 1. For each scenario, we generate 1,000 replicates and we estimate the ML values for the three models above and a model of independence with two parameters *μ*_1_ and *μ*_2_ (*H*_0_). We then compute the LRTs between four pairs of nested submodels (Figure 4a). LRT_1_ and LRT_4_ test whether adding a single parameter 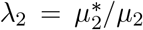 significantly improves the likelihood, demonstrating an induction of *E*_1_ on *E*_2_. LRT_2_ and LRT_3_ test whether the addition of two parameters (*v*_1_ and *v*_2_) significantly improves the likelihood, demonstrating that rate changes occur regardless of the interactions. When none of the LRTs is significant, a scenario of no induction+no loss (referred to as *H*_0_) is inferred. When LRT_1_ and LRT_4_ are the only two significant ones, a scenario of induction+no loss is inferred. When LRT_2_ and LRT_3_ are the only two significant ones, a scenario of loss+no-induction is inferred. When three LRTs are significant, a scenario of induction+loss is likely. Finally when all LRTs are significant, a scenario of induction+loss is inferred.

**Table 1:** Rates used for the simulations.

| | $\mu_1$ | $\mu_1^*$ | $\mu_2$ | $\mu_2^*$ | $\nu$ |
| --- | --- | --- | --- | --- | --- |
| Induction | 10 | 5 | 10 | [5, 5000] | $\mu$ |
| Loss | [5, 5000] | $\mu_1$ | 5 | 5 | 0 |
| Induction+Loss | 25 | 5 | 25 | [5, 5000] | 0 |

## Analysis of correlated evolution between Watson-Crick pairs

We applied our method to detect correlated evolution between Watson-Crick pairs of nucleotides in bacterial sequences of ribosomal ribonucleic acid sequences (rRNA). We downloaded 55 sequences of rRNA 16S from the SILVA rRNA database (Pruesse et al., 2007; Quast et al., 2013). Of those, 54 are sequences of gamma-enterobacteria and one from a beta-enterobacteria, used as an outgroup. Next, we aligned the sequences with Muscle v3.6 (Edgar, 2004) and trimmed all poorly aligned positions using Gblocks v0.91b with the default parameter values (Castresana, 2000; Talavera and Castresana, 2007), resulting in 1,233 aligned nucleotides. We next inferred a phylogenetic tree using PhyML (Guindon et al., 2010) with a generalized time reversible substitution model. Finally, we inferred the ancestral states by ML with BayesTraits Multistate (Pagel et al., 2004). We assimilated any substitution on any branch of the tree as an evolutionary event, disregarding the exact nature of the substitution. The list of the 55 species, the filtered alignment and the input tree (with or without events) are provided as Supplementary Material (see also Figure S1 for the displayed phylogenetic tree).

We evaluate the performance of two tests for independence. LRT_23_ compares the ML estimate of a 2-parameter model with no induction (*μ*_1_ and *μ*_2_) to a 3-parameter model with reciprocal induction (*μ*_1_, *μ*_2_ and 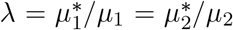). On the other hand, LRT_45_ compares the ML estimate of a 4-parameter model of indepence (*μ*_1_, *μ*_2_,*v*_1_ and *v*_2_) to a 5-parameter model with reciprocal induction (*μ*_1_, *μ*_2_, *v*_1_, *v*_2_ and 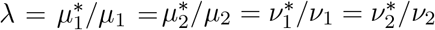).

To compare our method to state-of-the-art tools to detect correlated evolution, we selected four additional methods to detect correlated evolution in the rRNA alignment: two methods based on counts of co-occurrences (epics-Id (Behdenna et al., 2016) and CoMap (Dutheil et al., 2005)), one method based on Mutual Information corrected for the phylogeny (MIp (Gloor et al., 2005)) and the ML method BTDiscrete. To compute the correlated evolution with BTDiscrete, the current available implementation allows to test only binary states. Therefore, in the alignment positions where we observed at least three states, we replaced the least represented nucleotide(s) by a gap (i.e., considered as undefined character in the program).

For each of the tools, we ranked the 75 pairs that yielded the lowest p-values. To compute the distance in Å between each pair of nucleotides, we extracted the 3D position of the nucleotides in the crystal structure of the 16S ribosome (Korostelev et al., 2006) and used the barycenter as the reference position of each nucleotide.

### Implementation

The inference program, *epocs*, has been implemented in C language and is available at http://bioinfo.mnhn.fr/abi/public/EpoCs/. Simulations were performed using a home made simulator written in Ocaml language, available at the same url.

## Results

### Simulated data

We first assessed the power of the method on simulated data. Simulations give an overview of the theoretical power and the limitations of the method. Through different controlled scenarios, we evaluate (a) to what extent the method correctly estimates the occurrence rates, (b) its power to reject independence and (c) its ability to find the strength and nature of the interactions between the two evolutionary processes.

#### Estimating the occurrence rates

We first tested the accuracy of the method to retrieve arbitrarily selected occurrence rates (*μ*_1_ = 5, *μ*_2_ = 4, 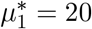 and 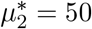) for 10,000 replicates. We observed that the distribution of all four rates shows a peak typically at the true values, demonstrating that our framework is indeed able to correctly estimate the rates (Figure 2). Notably, the basal rates are estimated with less bias (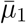 = 4.8 (sd = 2.4) and 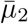 = 3.9 (sd = 2.1)) compared to the excited rates (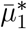 = 28.4 (sd = 58.2) and 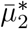 = 80.1 (sd = 74.0)). The dispersion of estimates around the true values mostly comes from the stochasticity of the simulations. Indeed, since the events are distributed according to Poisson processes, in some cases we can get more or fewer occurrences than expected. This also explains why we observe peaks at low rates that correspond to one or two occurrences. As the current implementation does not estimate rates larger than 1000, the last bin (1000+) includes all large rates.

**Figure 2:**
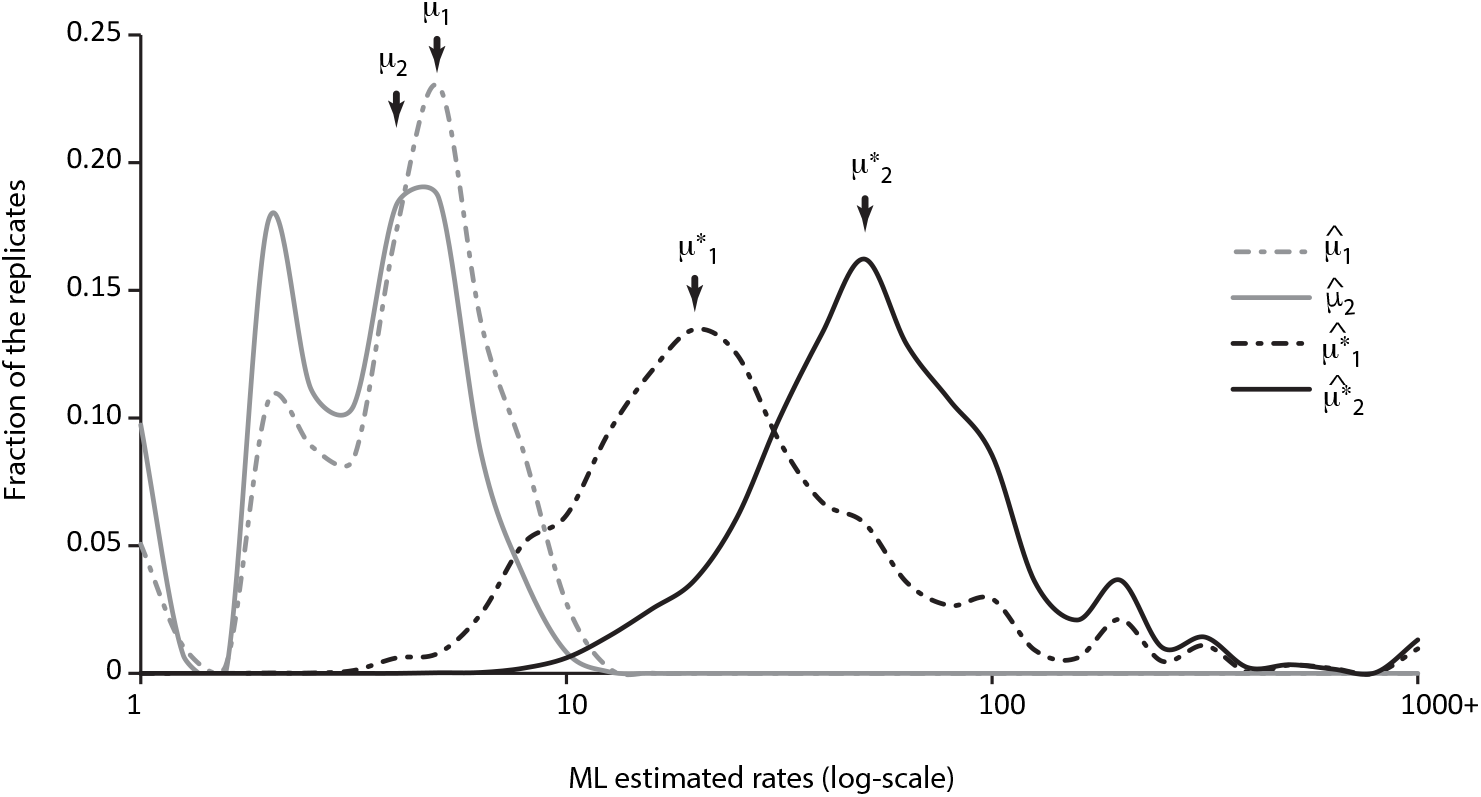
Empirical distribution of 10,000 replicates of rate estimation by maximum likelihood. For each replicate, we simulated on a perfectly symmetric tree with 64 leaves, occurrences of events with the following rates: *μ*_1_ = 5, *μ*_2_ = 4, 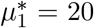 and 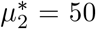. The different peaks at low rates correspond to cases where the numbers of realized events are small (1 or 2).

In addition, we found that the estimation of the lambda parameters approaches the expected values, where the estimation of the smaller *λ*_1_ is more accurate than the estimation of *λ*_2_ (Figure S2).

#### Power to detect induction

We further assessed the power of LRTs to detect an induction from *E*_1_ on *E*_2_ and compared them to the power of our previously published method based on counts in the phylogeny (Behdenna et al., 2016) and the ML method implemented in BayesTraits Discrete (Pagel, 1994). The power is estimated by the fraction of replicates that reject significantly the null model of independence with a p-value lower than 0.05.

The first simulated scenario is an asymmetric induction of *E*_1_ on *E*_2_ with three distinct parameters: *μ*_1_, *μ*_2_ and 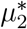 and an induction rate 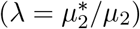 varying from 1 (no induction) to 1000 (very strong induction); all *v* values are equal to the corresponding μ.

The two computed LRTs, one comparing an independence model with two parameters and the simulated model, and the second comparing the independence model to an overparametrized model, outperform our previous method and the BTDiscrete module (Figure 3a). In particular, for inductions of *λ* > 10, LRTs show a very good power (above 60%) that reaches 98% for inductions *λ* > 100. Furthermore, we observed that the LRT based on the overparametrized model is slightly less powerful than the LRT based on the simulated model, suggesting that our method is capable of identifying the correct mode of correlated evolution.

**Figure 3:**
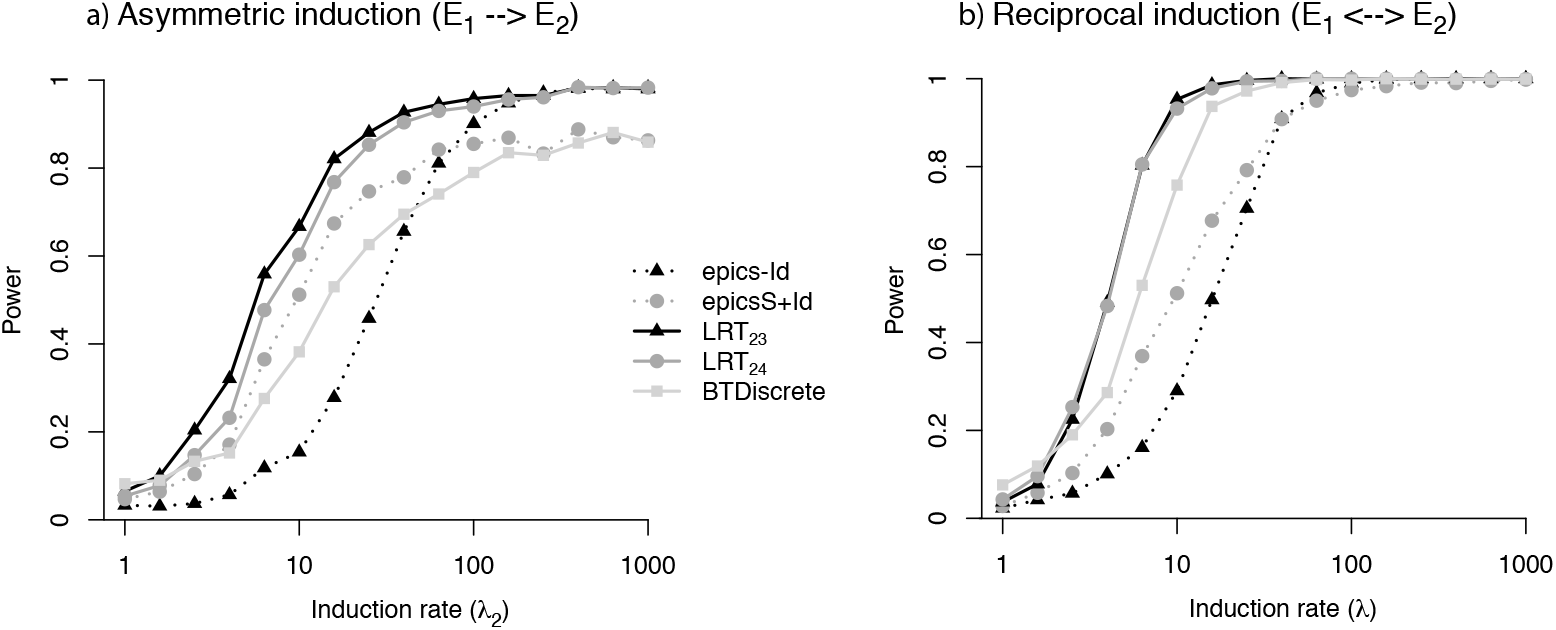
Power of likelihood-ratio tests. We compare the power of epocs to the power of our previous method, which computes exact p-values based on counts of cooccurring events (epics-Id) or co-occurring and sequentially ordered events (epicsS+Id). We also compute the result of BayesTraits Discrete (BTDiscrete). Increasing lambda values represent greater induction of *E*_1_ events on *E*_2_ events. Two LRTs are computed: LRT_23_ correspond to the LRT between *H*_0_ (2-parameter model) to *H*_1_ (3-parameter model, identical to the parameters used in the simulations), and LRT_24_ correspond to the LRT between *H*_0_ and *H*_2_ (overparametrized 4-parameter model). Thresholds of the LRT are computed assuming a *χ*^2^ distribution with one or two degrees of freedom accordingly. We retained p-values that are lower than 5% (a) Power to detect correlated evolution on a simulated scenario of induction with the three parameters *μ*_1_ = 5, *μ*_2_ = 5 and 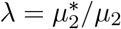, increasing from 1 to 1000. (b) Power to detect correlated evolution on a simulated scenario of reciprocal induction tuned by 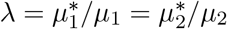, increasing from 1 to 1000.

The second scenario is a symmetric (reciprocal) induction between *E*_1_ and *E*_2_ with three distinct parameters: *μ*_1_, *μ*_2_ and 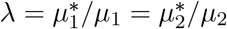 and *λ* ∈ [1,1000]. Similarly to the one-way induction, both LRTs outperform our previous method and the BTDiscrete module (Figure 3b). In this case of reciprocal induction, the power of LRTs reaches 95% already for inductions *λ* > 10.

#### Power to detect induction and/or irreversible loss

We next assessed the power of LRTs to discriminate the effect of *interactions between the events* (modeled by the induced rates *μ\**) and *intrinsic rate changes* (modeled by the *v* rates). More specifically, we designed three different scenarios where E_1_ events precede *E*_2_ events: (a) a scenario of induction, where E_1_ events favors *E*_2_ events, (b) a scenario of irreversible loss, where *E*_1_ events have a higher rate of occurrence and both events can occur only once, and (c) a mixed scenario of asymmetric induction and irreversible loss.

The exact rates that we used in the simulations are indicated in Table 1.

To assess the presence of induction and/or loss, we computed four LRTs to compare four pairs of nested models (Figure 4a). In particular, we aim to demonstrate that our method correctly identify the likely scenario underpinning the evolution of the simulated traits: a scenario of induction, a scenario of loss, or both.

**Figure 4:**
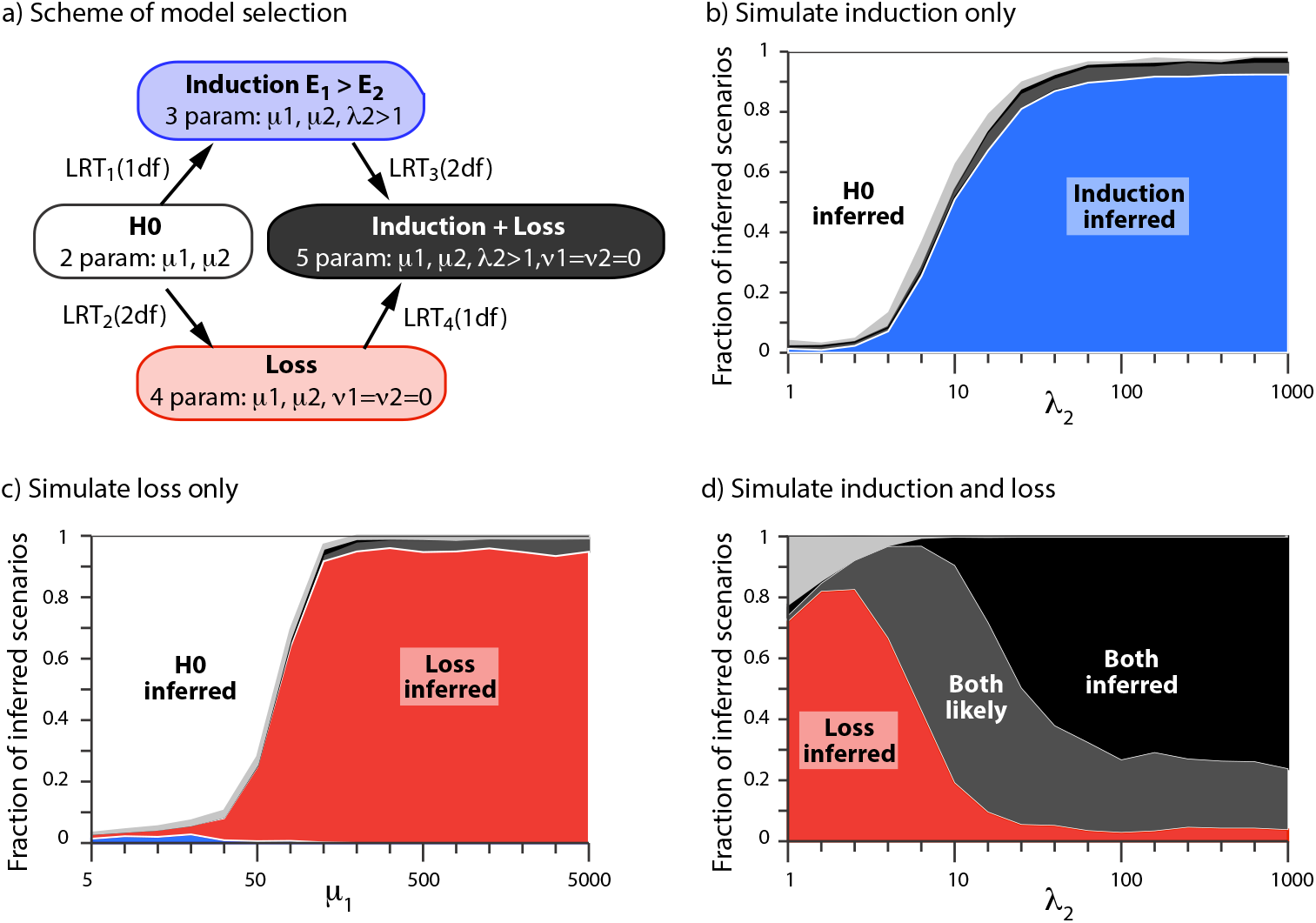
Power to assess correctly whether there is an induction, a loss or both. (a) Model selection and color code. All LRTs compare nested models with at least two rates (*μ*_1_ and *μ*_2_) and optional induction 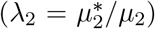 and/or loss (*v*_1_ = *v*_2_ = 0). Induction is inferred (blue area) when LRT_1_ and LRT_4_ are the only significant tests. Loss is inferred (red area) when LRT_2_ and LRT_3_ are the only significant tests. Both induction and loss are likely (dark grey area) when LRT_1_ and LRT_2_ are significant as well as either LRT_3_ or LRT_4_. Both are inferred (black areas) when all four tests are significant. White areas mark cases where no LRT is significant. All other cases are represented by light grey areas. (b-d) Results for the simulated scenarios. For each value of the x-axis, we simulated 1,000 replicates of one scenario. The y-axis indicates the fraction of each scenario that is inferred. (b) Scenario of induction of *E*_1_ on *E*_2_, with *μ*_1_ = *μ*_2_ = 10 and for increasing values of *λ*_2_; (c) Loss scenario, where *v*_1_ = *v*_2_ = 0, for increasing values of *μ*_1_; (d) Induction+loss scenario, with *μ*_1_ = *μ*_2_ = 25 and for increasing values of *λ*_2_.

In a simulated scenario of asymmetric induction, the model of induction and no loss is correctly preferred, provided that *λ*_2_ is large enough (*i.e.*, *λ*_2_ > 10) (Figure 4b). In a simulated scenario of loss and no induction, the correct scenario is inferred, provided that *μ*_1_ is large enough (Figure 4c). In other words, *E*_1_ events must occur rapidly and only once (followed by *E*_2_ events, that occur also once but later). Finally, we simulated a case with irreversible loss and induction. Evidence supporting both rate changes (intrinsic plus interaction) is higher as the induction of *E*_1_ on *E*_2_ gets larger (Figure 4d). When the induction is not very strong, the method only perceives the effect of loss.

From this set of simulations, we conclude that model selection based on LRTs can be used to assess the presence of interactions between the two types of events as well as intrinsic rate changes that are independent of any interaction.

### Analysis of rRNA 16S nucleotides

The ribosomal ribonucleic acid (rRNA) is a major component of the ribosome. The rRNA sequences are very well conserved, up until the last universal common ancestor. Hence, they have been used to show the diversity of living organisms, in particular the existence of the three kingdoms of life (Fox and Woese, 1975; Woese and Fox, 1977). The RNA 3D structure has been resolved experimentally (Doty et al., 1959; Leontis and Westhof, 2001), and its folding reveals physical interactions between nucleotides. In particular, the single-stranded structure of rRNA is in a large part determined by Watson-Crick pairs (hereafter WC pairs) that form stems. As the rRNA structure is strongly stabilized by stems of WC pairs, the mutation of one nucleotide involved in a WC pair is frequently deleterious and is typically only observed together with a second compensatory mutation on the paired nucleotide. WC pairs constitute a classical case of correlated evolution at the molecular level (Chiu and Kolodziejczak, 1991; Leontis and Westhof, 1998; Moore, 1999). Since both nucleotides of a WC pair are in strong interaction, we expect a very short time lag, if any, between the two mutations at paired bases.

#### WC pairs show evidence of correlated evolution

We used rRNA WC pairs as a positive control to evaluate the performance of two tests for independence: LRT_23_ compares the ML estimate of a 2-parameter model with no induction to a 3-parameter model with reciprocal induction. LRT_45_ compares the ML estimate of a 4-parameter model with no induction to a 5-parameter model with reciprocal induction. The rationale is to verify whether including switches of intrinsic rates improves the performance of LRTs. We also report the performance of our previous method based on counts of co-occurring events (epics-Id) and three additional methods to detect correlated evolution. The comparison between the pairs that are significantly coevolving and the documented WC pairs shows an overall good overlap between them (Table 2). The stronger the evidence of correlated evolution *(i.e.,* the smaller the p-values), the higher the enrichment of WC pairs. We observed that adding variable intrinsic rates (allowing *v* rates to differ from *μ* rates) improves the specificity of LRTs (at the cost of only a marginal loss of sensitivity), so we only used the LRT_45_ in the following and simply refer to it as LRT. The lower performance of BTDiscrete might stem from partially masking alignment positions with at least three different nucleotides (i.e., only the two most frequent alleles are considered in such positions). We found 34 WC pairs in these portions, hence decreasing the inference of correlated evolution for possible pairs.

**Table 2:** Summary of the rRNA nucleotide pairs that evolve in a correlated manner for different risk values. For each method, N indicates the number of pairs with evidence of correlated evolution for the risk given in the first column; WC is the number of Watson-Crick pairs among them.

| p-value | epics-Id |  | LRT<sub>23</sub> |  | LRT<sub>45</sub> |  | CoMap |  | Mip |  | BTDiscrete |  |
| --- | --- | --- | --- | --- | --- | --- | --- | --- | --- | --- | --- | --- |
|  | N | WC | N | WC | N | WC | N | WC | N | WC | N | WC |
| 0.05 | 3,006 | 74 | 9,464 | 89 | 6,850 | 86 | 6,519 | 87 | 2,219 | 83 | 1,012 | 63 |
| $10^{-2}$ | 698 | 57 | 2,940 | 74 | 2,477 | 75 | 2,098 | 83 | 202 | 51 | 186 | 47 |
| $10^{-3}$ | 101 | 40 | 752 | 61 | 426 | 59 | 421 | 60 | 50 | 34 | 31 | 23 |
| $10^{-4}$ | 39 | 29 | 266 | 50 | 118 | 45 | 118 | 39 | 28 | 25 | 8 | 8 |
| $10^{-5}$ | 22 | 20 | 68 | 39 | 49 | 37 | 40 | 31 | 7 | 7 | 3 | 3 |
| $10^{-6}$ | 12 | 11 | 37 | 34 | 30 | 28 | 0 | 0 | 0 | 0 | 0 | 0 |

However, we note that a large fraction of the 477 WC pairs show no evidence of correlated evolution, regardless of the method. At least two reasons can be put forward: the alignment filtering procedure and the monomorphic positions in the alignment. Indeed, out of the complete set of 477 WC pairs, only 349 WC pairs (0.05% of the 759,528 unmasked possible nucleotide pairs) have both their positions unmasked after Gblocks. Furthermore, the alignment contains 422 polymorphic sites, reducing the dataset to a maximum of 106 WC pairs that could be detected among a total of 88,831 polymorphic possible nucleotide pairs. Most of the 106 WC pairs show evidence of correlated evolution for a risk of 5%, but only ~ 30 very strong evidence of correlated evolution (p-value < 10^-6^).

To better understand what type of WC pairs show a significant pattern of correlated evolution, we computed the induction rate *λ* of all 106 WC pairs, regardless of the significance of the LRT (Figure 5). Remarkably, we observed that the 75 WC pairs with decent support for correlated evolution (p-value < 1%) have stronger induction rate than the others. Any WC pair with an induction rate of *λ* > 100 has significant support for correlated evolution at this risk.

**Figure 5:**
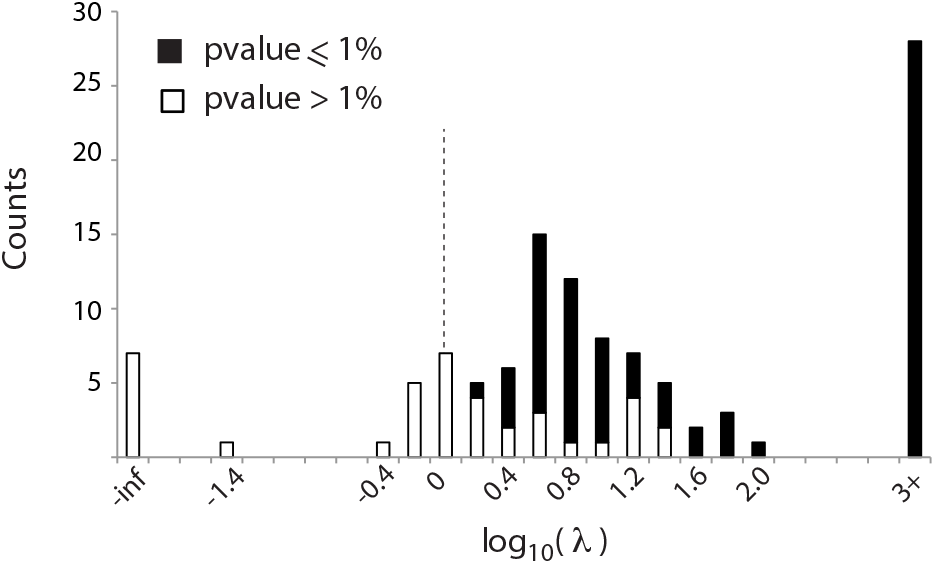
Induction rates of Watson-Crick pairs in 16S rRNA sequences of 54 enterobacteria. Assuming a model with five parameters (*μ*_1_, *μ*_2_, *v*_1_, *v*_2_ and 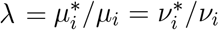), we estimated the reciprocal induction rate of mutation of one site onto the other. The rates are colored in black if there is statistical support for correlated evolution (at a risk of 1%) from the comparison between the model with five parameters to the model of independence with four parameters.

#### What other nucleotide pairs show evidence for correlated evolution?

We selected three additional methods to infer correlated evolution and compared the amount of WC pairs detected by each tool. For each method, we ranked the 75 best nucleotide pairs, *i.e.,* the pairs that show the strongest statistical evidence for correlated evolution (disregarding the significance of p-values). We then used the 3D positions of the nucleotides in the crystal structure of the 16S ribosome to compute the distance between each pair of nucleotides. Each pair was then tagged depending on whether it was a WC pair, a non-WC pair with both nucleotides close in space (< 10^Å^) or a distant pair. We found that the LRT method is marginally more specific at detecting WC pairs (Figure 6a). In addition, epics-Id was also able to detect significant correlated evolution among close nucleotides that are not WC pairs.

**Figure 6:**
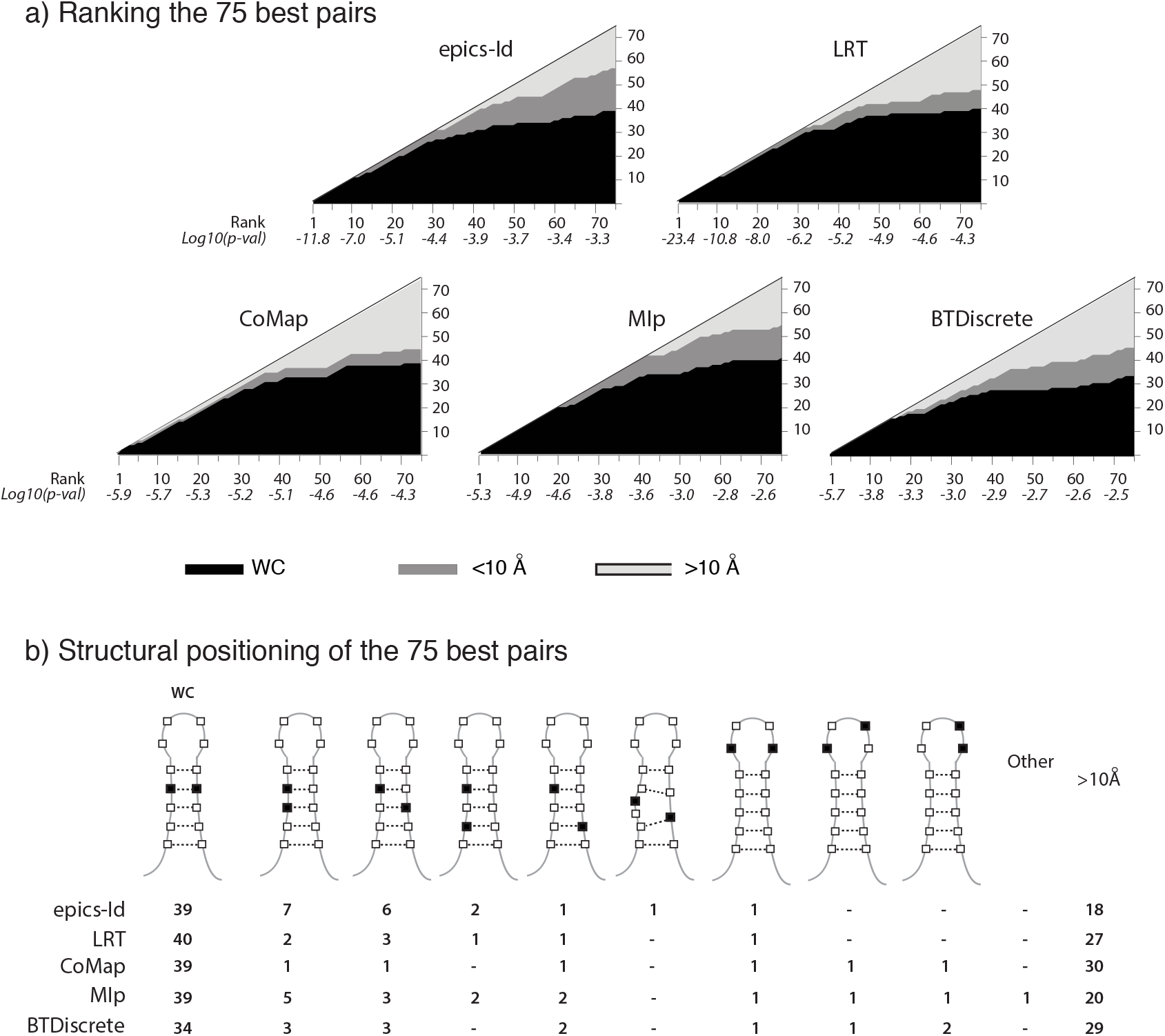
Analysis of the 75 best nucleotide pairs of 16S showing correlated evolution. (a) We report the nature of the 75 best pairs (the ones with the lowest probabilities) among all nucleotide pairs, according to five different methods: our previous method with co-occurrences (epics-Id), the LRT based method (see legend of Figure 5) using our likelihood framework, the CoMap method based on co-occurrences on the tree, the MIp method based on Mutual Information, corrected for phylogeny, and the BTDiscrete module. Pairs are colored in black when they are Watson-Crick, on dark grey when they are closer than ten ^Å^ in the 3D structure and in light grey otherwise. Below their rank, we also report their associated p-value. (b) Relative structural positions on the stems-loops for the nucleotides of the 75 best pairs, according to the five methods.

To further characterize the non-WC nucleotide pairs that show patterns of correlated evolution, we mapped the nucleotide positions on the rRNA secondary structure: stems of WC pairs and associated loops. Several non-WC pairs close in space belong to the same stem and are either consecutive (positions *k* and *k* + 1, i.e., stacked nucleotides) or shifted by one when compared to WC pairs (Figure 6b). This suggests that alternative non-WC interactions also constrain the evolution of the 16S rRNA.

## Discussion

In this study, we have developed a likelihood framework based on a minimal model of correlated evolution between two evolutionary processes that generate discrete events on a given phylogenetic tree. This model depends on at most eight parameters, four for each process, that represent *interactions* between the two processes (the starred rates) and intrinsic rate switches (*v* ≠ *μ*). For a given tree with mapped evolutionary events, we estimate the parameters by maximum likelihood for any (sub)model defined with (a subset of) the eight parameters. Maximizing this probability is equivalent to searching for an optimal set of parameters describing the processes that led to the observed distribution of the events on the tree. Nested models also permit the direct comparison of scenarios using likelihood ratio tests, in particular to assess the support for interaction and/or intrinsic rate switches.

Using simulated controlled scenarios, we have shown that our method is accurate for estimating the occurrence rates and has good power to assess the presence of interactions despite the use of limited data (symmetric trees with only 64 leaves). Moreover, we show that the method can assess correctly which process can lead to ordered pairs of events: differential rates with irreversible loss or interaction between the two processes. Finally, based on an analysis of nucleotide pairs in an alignment of 16S rRNA sequences sampled in 54 enterobacteria, we show that the method detects correlated evolution in many polymorphic WC pairs, especially among the ones that exhibit strong reciprocal induction. In addition, we show that several pairs that are not WC but in the same stem also show evidence of correlated evolution. The comparison of performance with a selection of four previously existing methods demonstrates that the current framework is more specific at detecting WC pairs and can additionally estimate the strength of their reciprocal induction.

As the model has a small number of parameters, the computation time of likelihood maximization is fast enough to analyze correlated evolution in a very large number of pairs. In the 16S analysis, we had to maximize 2,304,091 likelihood functions (finding all possible orders of events for all pairs). The total computation time on a 8Ghz laptop, without any parallelization, is 11 minutes for the 2-parameter model of independence (*μ*_1_ and *μ*_2_), 71 minutes for the 4-parameter model of independence, and at most 240 minutes for the full 8-parameter model (that was not used in the analysis above). This suggests that MCMC Bayesian surface exploration, that typically requires millions of likelihood estimations, could be run to analyze cases on a dataset of a reasonable size. Computation time remains, however, orders of magnitude larger than in the methods based on counts we have developed before (Behdenna et al., 2016), which takes only four seconds to compute correlated evolution from counts of co-occurrence.

The framework described here relates to the one proposed by Pagel (1994), as they both rely on likelihood computation of an explicit continuous time Markov chain. In both frameworks, the independence model is nested in a subspace of the more general model of correlated evolution, allowing to use statistical tests such as LRT to assess the statistical support for independence. However, our minimal framework differs from the one of Pagel (1994) in several important aspects. On of the major difference is that our model has at most eight parameters, whereas Pagel (1994) computes the transition rates between all possible pairs of states for the two traits. In the case of two binary traits, Pagel (1994) has eight parameters but it has hundreds for sites with 20 amino acids.

If we restrict our minimal framework to the 4-parameter model of independence (i.e., the null model of Pagel (1994)) for binary traits, and the largest 8-parameter model (i.e., the full model of Pagel (1994)), both frameworks are similar but not identical. Indeed, our model has twelve hidden states compared to four for Pagel (1994) and there is no bidirectional mapping between the transition rates.

However, our minimal model is flexible enough that one can test for likelihood improvement due to the addition of intrinsic rates changes (*v* ≠ *μ*) and/or addition of interaction between the processes (*μ*^*^ = *μ*). Moreover, the time to maximize the likelihood for BTDiscrete in the Watson-Crick analysis (240 minutes to compute 83,028 pairwise site comparisons) is much higher than our implementation.

More generally, our minimal model constitutes a fast and powerful tool to assess the statistical support for any scenario that is defined within the framework, ignoring extra unnecessary parameters. In addition, the comparison of the various rates of our minimal model is straightforward as they have an explicit biological interpretation (e.g., the *λ* values immediately categorizes the interaction as an induction or a repression).

A drawback of our method is that it is not designed to infer correlated evolution for processes for which occurrence rates are tuned by more than two states, e.g., when multiple amino acid transitions occur with different rates in the phylogenetic tree at the same sites. Furthermore, as the method relies on a phylogenetic tree with mapped evolutionary events, it will suffer from uncertainties in phylogenetic inference and ancestral state reconstruction. Indeed, the method exploits the branch lengths to estimate the rates of the evolutionary processes on the tree. As such, trees estimated on very close sequences or on very divergent ones should be handled with care. In addition, since deep branches are particularly subject to uncertainties, one has to be careful with the tree used as input. Furthermore, our method assumes that evolutionary events are correctly placed on the tree. We therefore recommend great care in the reconstruction of the ancestral states. When possible, likelihood-based methods, such as ML (Ishikawa et al., 2019; Pupko et al., 2000) or Bayesian methods (Bouckaert et al., 2014; Pagel et al., 2004) should be preferred over parsimony. Nonetheless, we note that when there are only few events on the tree, all methods will likely perform similarly. One possible extension of our framework would be to integrate over all possible ancestral reconstructions. Altogether, we suggest that the method presented here is best suited for phylogenies that are not too deep and where the evolutionary events of interest are sparsely located.

One interesting feature of our framework is its inherent flexibility that allows the analysis of diverse types of data. Indeed, any discrete evolutionary event, regardless of its nature, that is mapped on a phylogenetic tree can be analyzed, whether it is molecular (e.g., substitution) or non-molecular (e.g., gain or loss of a character or of a biological function).

As a concluding remark, we would like to mention that genome-wide association studies (GWAS) can be seen as computing correlated evolution between genomic variants and a phenotypic trait of interest (Achaz and Dutheil, 2021). As any method of correlated evolution, GWAS also suffer from phylogenetic inertia due to stratification of the population into subpopulations (Price et al., 2006).

For the special case of bacteria, the TreeWAS framework was recently developed to perform GWAS-like analyses on a phylogenetic tree (Collins and Didelot, 2018). In short, the TreeWAS method performs an association analysis between genetic variants and the phenotype of interest, both conditioned on a phylogenetic tree. This suggests that our framework presented here, that can assess and quantify correlated evolution between discrete traits mapped on a phylogeny, can also be used to perform studies similar to TreeWAS.

## Acknowledgments

The authors want to thank EPC Rocha and R Kulathinal for their critical reading and helpful comments on the manuscript as well as O Tenaillon for fruitful discussions on the work. This work was supported by a state grant from the *Agence Nationale de la Recherche* through the *Investissements d’Avenir* program ANR-16-CONV-0005. AB, MG, PP, AL and GA thank the Center for Interdisciplinary Research in Biology (CIRB, College de France) for funding. The authors have declared that no competing interests exist.

## Appendix

### A Initial and final states

The following tables sum up the final rate state corresponding to each initial rate state, for each possible case. Note that the states where both events have a starred occurrence rate are forbidden.

#### No event on the branch

In this particular case, the final state is always the same as the initial state.

#### One occurrence of *E_i_* only on the branch

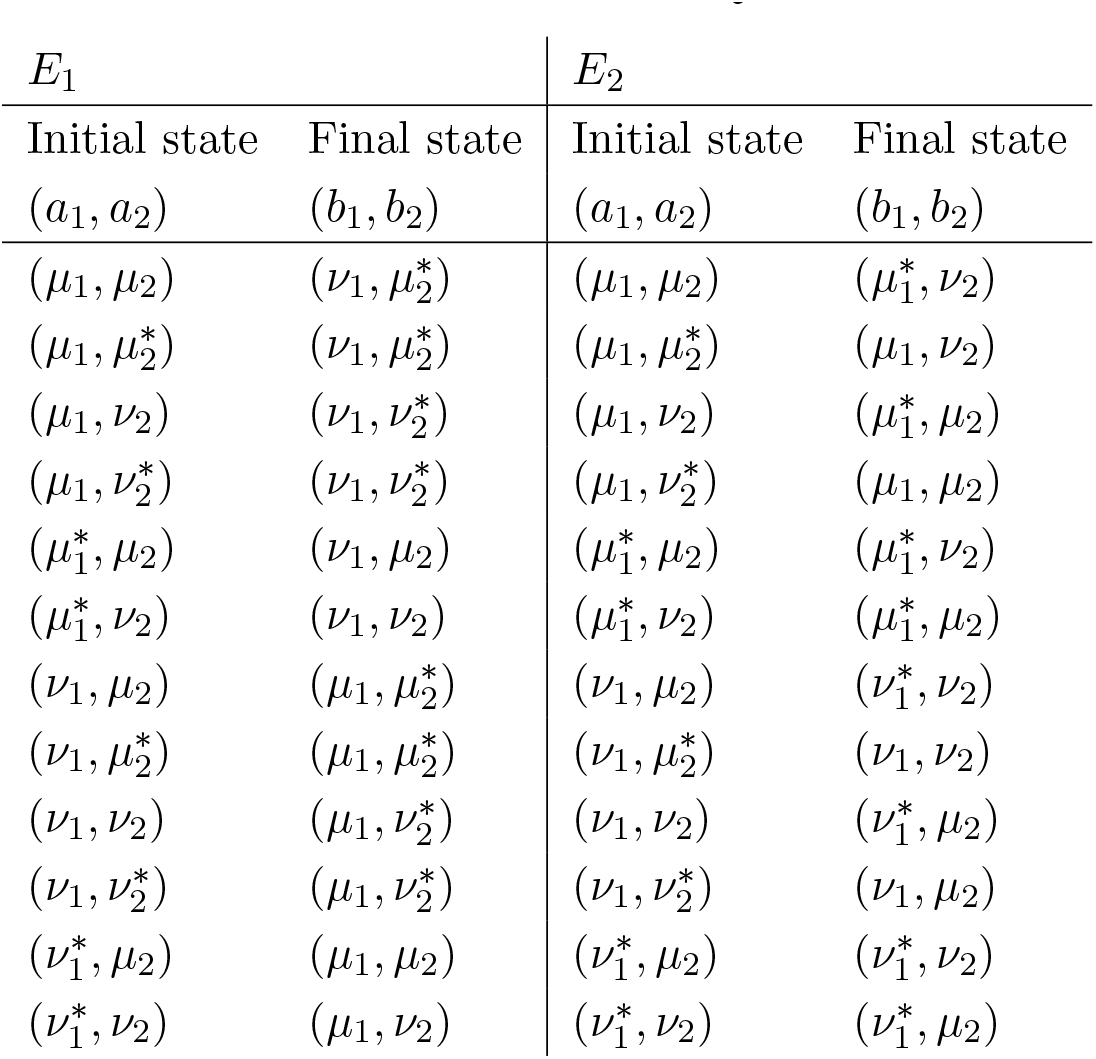

#### Both events are present on the branch

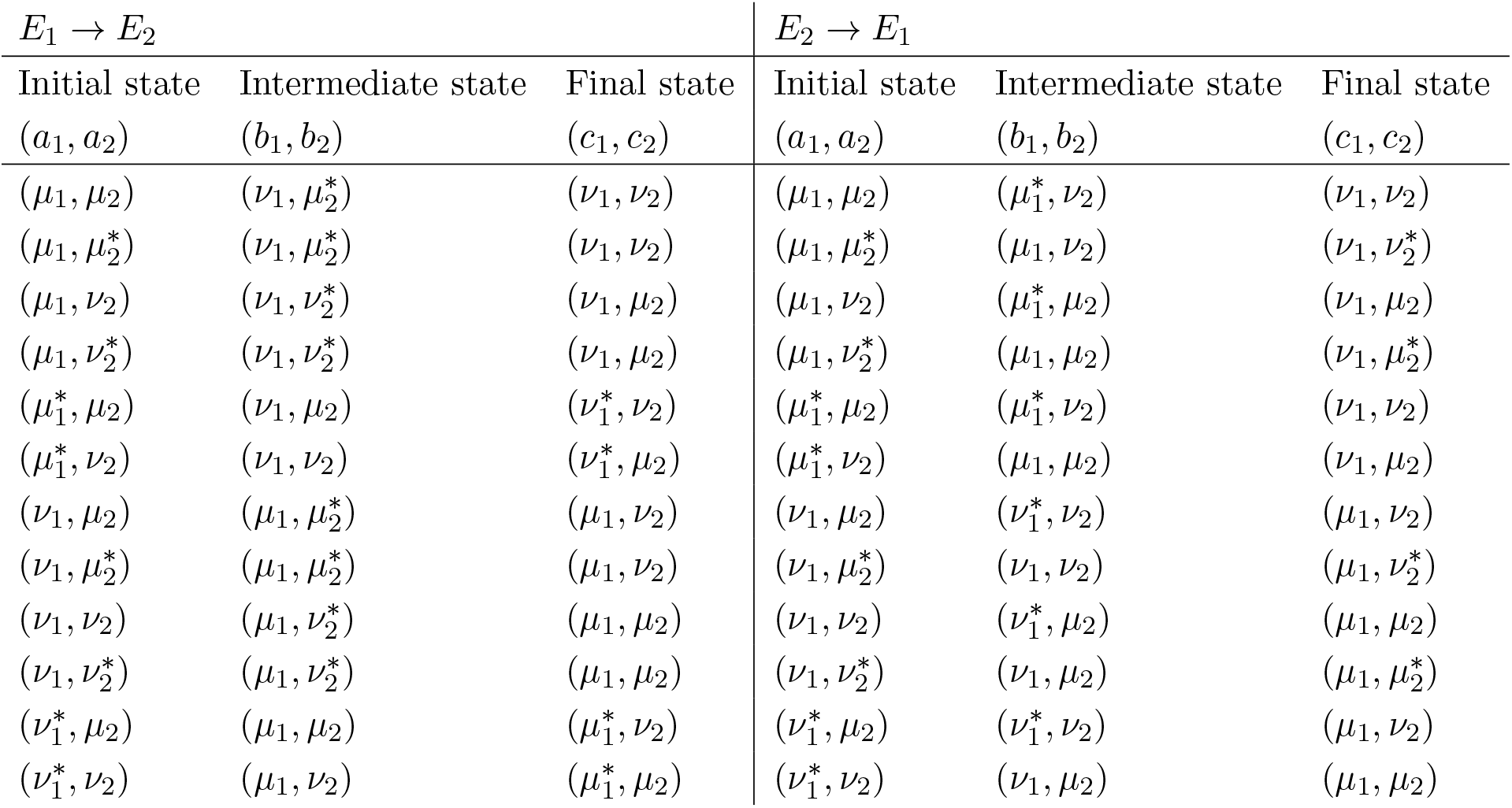

### B Likelihood function for a single branch

Let *T* be a tree, and 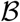 a branch of this tree, of length *l*. We consider two events *E_i_*, *i* ∈ {1,2}, of respective occurrence rates (*μ*_1_, 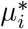,*v*_1_, 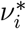). Each event is assumed to occur at most once on 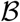.

We detail here the likelihood function for this single branch, the three possible cases being treated separately. Let 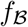 be the likelihood for the branch 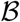, given the parameter values.

#### No event on the branch

Let *a*_1_ (resp. *a*_2_) be the occurrence rate for *E*_1_ (resp. *E*_2_) on the branch.

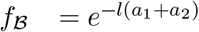

#### One occurrence of *E_i_*, *i* ∈ {1, 2} on the branch

In this case, we define:

- *a*_1_ (resp. *a*_2_) as the occurrence rate for *E*_1_ (resp. *E*_2_) before the occurrence of *E_i_* on the branch
- *b*_1_ (resp. *b*_2_) as the occurrence rate for *E*_1_ (resp. *E*_2_) after the occurrence of *E_i_* on the branch.

Those parameters take different values depending on the initial states on the branch, and according to the model. We easily get

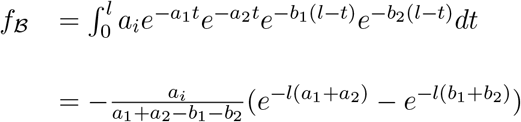

When (*a*_1_ + *a*_2_ − *b*_1_ — *b*_2_) → 0:

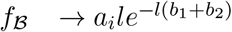

#### One occurrence of each event on the branch

In this subsection, we assume that the occurrence of *E*_1_ precedes the occurrence of *E_j_* on branch 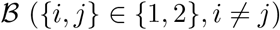.

By analogy with the previous case, we define:

- *a*_1_ (resp. *a*_2_) as the occurrence rate for *E*_1_ (resp. *E*_2_) before the occurrence of *E_i_* on the branch
- *b*_1_ (resp. *b*_2_) as the occurrence rate for *E*_1_ (resp. *E*_2_) after the occurrence of *E_i_* and before the occurrence of *E_j_* on the branch.
- *c*_1_ (resp. *c*_2_) as the occurrence rate for *E*_1_ (resp. *E*_2_) after the occurrence of *E_j_* on the branch.

Those parameters take different values depending on the initial states on the branch, and according to the model. We easily get

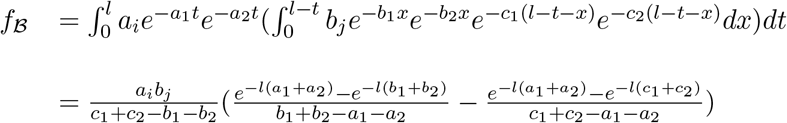

If all three denominators tend to 0,

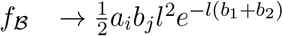

When *c*_1_ + *c*_2_ — *b*_1_ — *b*_2_ → 0,

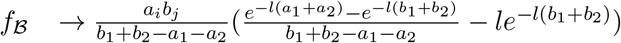

When *b_1_* + *b*_2_ — *a_i_* — *a*_2_ → 0,

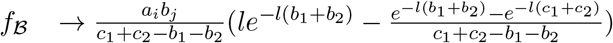

When *a_1_* + *a*_2_ — *c*_i_ — *c*_2_ → 0,

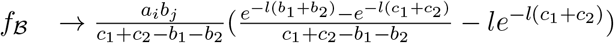

## Notes

https://doi.org/10.5061/dryad.dbrv15dzq

### Competing Interest Statement

The authors have declared no competing interest.

## References

Achaz, G. and J. Dutheil. 2021. Correlated evolution: models and methods. arXiv:2103.11809 [q-bio.PE].

Achaz, G., A. Rodriguez-Verdugo, B. S. Gaut, and O. Tenaillon. 2014. The reproducibility of adaptation in the light of experimental evolution with whole genome sequencing. Adv Exp Med Biol 781:211–31.

Baldassi, C., M. Zamparo, C. Feinauer, A. Procaccini, R. Zecchina, M. Weigt, and A. Pagnani. 2014. Fast and accurate multivariate Gaussian modeling of protein families: predicting residue contacts and protein-interaction partners. PLOS ONE 9:e92721.

Bateson, W. 1909. Heredity and variation in modern lights. chap. III, Pages 87–110 in Danvinand Modmz Science (A. Seward, ed.) gutenberg projet http://www.gutenberg.org/files/22430/22430-h/22430-h.htm ed. Cambridge University Press.

Baum, D. A. and M. J. Donoghue. 2001. A likelihood framework for the phylogenetic analysis of adaptation. Pages 24–44 in Adaptation and Optimality (S. Orzack and E. Sober, eds.). Cambridge University Press, New York.

Behdenna, A., J. Pothier, S. S. Abby, A. Lambert, and G. Achaz. 2016. Testing for Independence between Evolutionary Processes. Syst Biol 65:812–823.

Bitbol, A.-F. 2018. Inferring interaction partners from protein sequences using mutual information. PLOS Comput Biol 14:e1006401.

Bouckaert, R., J. Heled, D. Kühnert, T. Vaughan, C.-H. Wu, D. Xie, M. A. Suchard, A. Rambaut, and A. J. Drummond. 2014. Beast 2: A software platform for bayesian evolutionary analysis. PLOS Comput Biol 10:1–6.

Castresana, J. 2000. Selection of conserved blocks from multiple alignments for their use in phylogenetic analysis. Mol Biol Evol 17:540–52.

Chiu, D. K. and T. Kolodziejczak. 1991. Inferring consensus structure from nucleic acid sequences. Comput Appl Biosci 7:347–352.

Collins, C. and X. Didelot. 2018. A phylogenetic method to perform genome-wide association studies in microbes that accounts for population structure and recombination. PLOS Comput Biol 14:e1005958.

Dib, L., D. Silvestro, and N. Salamin. 2014. Evolutionary footprint of coevolving positions in genes. Bioinformatics 30:1241–9.

Dobzhansky, T. 1934. Studies on hybrid sterility. Zeitschrift für Zellforschung und Mikroskopische Anatomie 21:169–223.

Doty, P., H. Boedtker, J. R. Fresco, R. Haselkorn, and M. Litt. 1959. Secondary structure in ribonucleic acids. Proc Natl Acad Sci U S A 45:482–99.

Dutheil, J. and N. Galtier. 2007. Detecting groups of coevolving positions in a molecule: a clustering approach. BMC Evol Biol 7:242.

Dutheil, J., T. Pupko, A. Jean-Marie, and N. Galtier. 2005. A model-based approach for detecting coevolving positions in a molecule. Mol Biol Evol 22:1919–28.

Edgar, R. C. 2004. Muscle: multiple sequence alignment with high accuracy and high throughput. Nucleic Acids Res 32:1792–7.

Ekeberg, M., C. Lövkvist, Y. Lan, M. Weigt, and E. Aurell. 2013. Improved contact prediction in proteins: using pseudolikelihoods to infer Potts models. Phys Rev E Stat Nonlin Soft Matter Phys 87:012707.

Felsenstein, J. 1981. Evolutionary trees from dna sequences: a maximum likelihood approach. J Mol Evol 17:368–76.

Felsenstein, J. 1985. Phylogenies and the comparative method. The American Naturalist 125:1–15.

Fox, G. E. and C. R. Woese. 1975. 5s rna secondary structure. Nature 256:505–7.

Fraser, H. B., A. E. Hirsh, D. P. Wall, and M. B. Eisen. 2004. Coevolution of gene expression among interacting proteins. Proc Natl Acad Sci U S A 101:9033–9038.

Gloor, G. B., L. C. Martin, L. M. Wahl, and S. D. Dunn. 2005. Mutual information in protein multiple sequence alignments reveals two classes of coevolving positions. Biochemistry 44:7156–65.

Guindon, S., J.-F. Dufayard, V. Lefort, M. Anisimova, W. Hordijk, and O. Gascuel. 2010. New algorithms and methods to estimate maximum-likelihood phylogenies: assessing the performance of phyml 3.0. Syst Biol 59:307–21.

Harvey, P. H. and M. D. Pagel. 1991. The Comparative Method In Evolutionary Biology. Oxford University Press, U.S.A., Oxford; New York.

Ishikawa, S. A., A. Zhukova, W. Iwasaki, and O. Gascuel. 2019. A Fast Likelihood Method to Reconstruct and Visualize Ancestral Scenarios. Mol Biol Evol 36:2069–2085.

Ives, A. R. and T. Garland, Jr. 2010. Phylogenetic Logistic Regression for Binary Dependent Variables. Syst Biol 59:9–26.

Kondrashov, A. S., S. Sunyaev, and F. A. Kondrashov. 2002. Dobzhansky-muller incompatibilities in protein evolution. Proc Natl Acad Sci U S A 99:14878–83.

Korostelev, A., S. Trakhanov, M. Laurberg, and H. F. Noller. 2006. Crystal structure of a 70s ribosome-trna complex reveals functional interactions and rearrangements. Cell 126:1065–1077.

Kryazhimskiy, S., J. Dushoff, G. A. Bazykin, and J. B. Plotkin. 2011. Prevalence of epistasis in the evolution of influenza A surface proteins. PLOS Genet 7:e1001301.

Kulathinal, R. J., B. R. Bettencourt, and D. L. Hartl. 2004. Compensated deleterious mutations in insect genomes. Science 306:1553–4.

Leontis, N. B. and E. Westhof. 1998. A common motif organizes the structure of multihelix loops in 16 s and 23 s ribosomal rnas. J Mol Biol 283:571–583.

Leontis, N. B. and E. Westhof. 2001. Geometric nomenclature and classification of rna base pairs. RNA 7:499–512.

Marks, D. S., L. J. Colwell, R. Sheridan, T. A. Hopf, A. Pagnani, R. Zecchina, and C. Sander. 2011. Protein 3D Structure Computed from Evolutionary Sequence Variation. PLOS ONE 6:e28766.

Martin, L. C., G. B. Gloor, S. D. Dunn, and L. M. Wahl. 2005. Using information theory to search for co-evolving residues in proteins. Bioinformatics 21:4116–24.

Milligan, B. 1994. Estimating evolutionary rates for discrete characters. Pages 299–311 in Models in phylogeny reconstruction (R. W. Scotland, D. J. Siebert, and D. M. Williams, eds.) vol. Systematics Association Special Volume Number 52. Clarendon, Oxford, UK.

Moore, P. B. 1999. Structural motifs in rna. Annu Rev Biochem 68:287–300.

Morcos, F., A. Pagnani, B. Lunt, A. Bertolino, D. S. Marks, C. Sander, R. Zecchina, J. N. Onuchic, T. Hwa, and M. Weigt. 2011. Direct-coupling analysis of residue coevolution captures native contacts across many protein families. Proc Natl Acad Sci U S A 108:E1293–1301.

Muller, H. 1942. Isolating mechanisms, evolution, and temperature. Biology Symposium 6:71–125.

Neyman, J. and E. Pearson. 1933. On the problems of the most efficient tests of statistical hypotheses. Philos Trans R Soc Pages 289–337.

Orr, H. A. 1996. Dobzhansky, bateson, and the genetics of speciation. Genetics 144:1331–5.

Pagel, M. 1994. Detecting correlated evolution on phylogenies: a general method for the comparative analysis of discrete characters. Proc R Soc B 255:37–45.

Pagel, M. and A. Meade. 2006. Bayesian analysis of correlated evolution of discrete characters by reversible-jump markov chain monte carlo. Am Nat 167:808–25.

Pagel, M., A. Meade, and D. Barker. 2004. Bayesian estimation of ancestral character states on phylogenies. Syst Biol 53:673–84.

Pensar, J., S. Puranen, B. Arnold, N. MacAlasdair, J. Kuronen, G. Tonkin-Hill, M. Pesonen, Y. Xu, A. Sipola, L. Sánchez-Busó, J. A. Lees, C. Chewapreecha, S. D. Bentley, S. R. Harris, J. Parkhill, N. J. Croucher, and J. Corander. 2019. Genome-wide epistasis and co-selection study using mutual information. Nucleic Acids Res 47:e112–e112.

Phillips, P. C. 2008. Epistasis-the essential role of gene interactions in the structure and evolution of genetic systems. Nat Rev Genet 9:855–67.

Poelwijk, F. J., D. J. Kiviet, D. M. Weinreich, and S. J. Tans. 2007. Empirical fitness landscapes reveal accessible evolutionary paths. Nature 445:383–6.

Pollock, D. D., W. R. Taylor, and N. Goldman. 1999. Coevolving protein residues: maximum likelihood identification and relationship to structure. J Mol Biol 287:187–98.

Price, A. L., N. J. Patterson, R. M. Plenge, M. E. Weinblatt, N. A. Shadick, and D. Reich. 2006. Principal components analysis corrects for stratification in genomewide association studies. Nat Genet 38:904–909.

Pruesse, E., C. Quast, K. Knittel, B. M. Fuchs, W. Ludwig, J. Peplies, and F. O. Glöckner. 2007. Silva: a comprehensive online resource for quality checked and aligned ribosomal rna sequence data compatible with arb. Nucleic Acids Res 35:7188–96.

Pupko, T., I. Pe, R. Shamir, and D. Graur. 2000. A Fast Algorithm for Joint Reconstruction of Ancestral Amino Acid Sequences. Mol Biol Evol 17:890–896.

Quast, C., E. Pruesse, P. Yilmaz, J. Gerken, T. Schweer, P. Yarza, J. Peplies, and F. O. Glöckner. 2013. The silva ribosomal rna gene database project: improved data processing and web-based tools. Nucleic Acids Res 41:D590–6.

Schoniger, M. and A. von Haeseler. 1994. A stochastic model for the evolution of autocorrelated dna sequences. Mol Phylogenet Evol 3:240–7.

Shindyalov, I. N., N. A. Kolchanov, and C. Sander. 1994. Can three-dimensional contacts in protein structures be predicted by analysis of correlated mutations? Protein Eng Des Sel 7:349–358.

Talavera, G. and J. Castresana. 2007. Improvement of phylogenies after removing divergent and ambiguously aligned blocks from protein sequence alignments. Syst Biol 56:564–77.

Tillier, E. R. M. and R. A. Collins. 1995. Neighbor joining and maximum likelihood with rna sequences: Adressing the interdependence of sites. Mol Biol Evol 12:7–15.

Tufféry, P. and P. Darlu. 2000. Exploring a phylogenetic approach for the detection of correlated substitutions in proteins. Mol Biol Evol 17:1753–1759.

Van Valen, L. 1973. A New Evolutionary Law. Evol Theory 1:1–30.

Visser, J. A. G. M. d. and J. Krug. 2014. Empirical fitness landscapes and the predictability of evolution. Nat Rev Genet 15:480–490.

Weigt, M., R. A. White, H. Szurmant, J. A. Hoch, and T. Hwa. 2009. Identification of direct residue contacts in protein-protein interaction by message passing. Proc Natl Acad Sci U S A 106:67–72.

Weinreich, D. M., R. A. Watson, and L. Chao. 2005. Perspective: Sign epistasis and genetic constraint on evolutionary trajectories. Evolution 59:1165–74.

Welch, J. J. 2004. Accumulating dobzhansky-muller incompatibilities: reconciling theory and data. Evolution 58:1145–56.

Woese, C. R. and G. E. Fox. 1977. Phylogenetic structure of the prokaryotic domain: the primary kingdoms. Proc Natl Acad Sci U S A 74:5088–90.

Wright, S. 1932. The roles of mutation, inbreeding, crossbreeding and selection in evolution. Pages 356–366 in Proceedings of the Sixth International Congress on Genetics vol. 1.

Yeang, C.-H., J. Darot, H. Noller, and D. Haussler. 2007. Detecting the coevolution of biosequences an example of rna interaction prediction. Mol Biol Evol 24:2119–31.

Yi, X. and A. M. Dean. 2019. Adaptive Landscapes in the Age of Synthetic Biology. Mol Biol Evol 36:890–907.

